# METAMVGL: a multi-view graph-based metagenomic contig binning algorithm by integrating assembly and paired-end graphs

**DOI:** 10.1101/2020.10.18.344697

**Authors:** Zhenmiao Zhang, Lu Zhang

## Abstract

**Motivation:** Due to the complexity of metagenomic community, *de novo* assembly on next generation sequencing data is commonly unable to produce microbial complete genomes. Metagenomic binning is a crucial task that could group the fragmented contigs into clusters based on their nucleotide compositions and read depths. These features work well on the long contigs, but are not stable for the short ones. Assembly and paired-end graphs can provide the connectedness between contigs, where the linked contigs have high chance to be derived from the same clusters.

**Results:** We developed METAMVGL, a multi-view graph-based metagenomic contig binning algorithm by integrating both assembly and paired-end graphs. It could strikingly rescue the short contigs and correct the binning errors from dead ends subgraphs. METAMVGL could learn the graphs’ weights automatically and predict the contig labels in a uniform multi-view label propagation framework. In the experiments, we observed METAMVGL significantly increased the high-confident edges in the combined graph and linked dead ends to the main graph. It also outperformed with many state-of-the-art binning methods, MaxBin2, MetaBAT2, MyCC, CONCOCT, SolidBin and Graphbin on the metagenomic sequencing from simulation, two mock communities and real *Sharon* data.

**Availability and implementation:** The software is available at https://github.com/ZhangZhenmiao/METAMVGL.

## 1 Introduction

During long-term genetic evolution, animals, including humans, have formed complex ecosystems of symbiotic relationships with diverse microbes. The gut microbiome is a community with the highest microbial density in the human body, including thousands of microbial species mixed in varying proportions and constituting a dynamic system. Most gut microbes are difficult to isolate and culture in vitro. Metagenomic sequencing is designed to directly sequence a mixture of microbes and explore microbial compositions and abundances by data post-processing.

Due to the paucity of high-quality microbial reference genomes, current pipelines commonly target single genes or species using species-specific markers (Li et al. (2014); Truong et al. (2015)). But novel microbes could be lost by alignment-based approaches. Metagenome assembly is a promising strategy to explore the novel species by concatenating the shortreads into long contigs, which are regarded as the pieces of strain genomes. The fragmented contigs are further grouped into strain-specific clusters, called contig binning. This strategy have been widely adopted to explore the novel microbes from the human gut microbiome (Almeida et al. (2019, 2020); Consortium et al. (2010); Forster et al. (2019); Nayfach et al. (2019); Pasolli et al. (2019); Poyet et al. (2019); Zou et al. (2019)).

Many state-of-the-art contig binning algorithms have been developed by considering contig nucleotide compositions (tetranucleotide frequencies (TNF), k-mer frequencies, marker genes, codon usage) and sequence depth. MaxBin2 (Wu et al. (2016)) uses Expectation–Maximization algorithm to maximize the probability of a contig belonging to local cluster centers using TNF and sequence depth. These two types of information are also used in MetaBAT2 (Kang et al. (2019)) to calculate contig similarities. MetaBAT2 constructs a graph with contigs as vertices and their similarities as edges’ weights, which is partitioned into groups by applying a modified label propagation algorithm. CONCOCT (Alneberg et al. (2014)) applies Gaussian mixture models for contig clustering based on k-mer frequencies and sequence depth across multiple samples. Besides considering TNF, MyCC (Lin and Liao (2016)) also aggregates the contigs with complementary marker genes by affinity propagation. BMC3C (Yu et al. (2018)) applies codon usage in the ensemble clustering. These methods can not deal with the short contigs (commonly<1kb), because they might lead to unstable nucleotide composition distributions and sequence depth. We observed a majority of the contigs (89.55%, Table S1 *Sharon* with metaSPAdes) in the assembly graph were shorter than 1kb, which would be dropped by binning algorithms.

To rescue those short contigs, Mallawaarachchi et al. (2020) developed Graphbin to label the short contigs and correct the potential binning errors by employing label propagation on the assembly graph. In principle, the assembly graph should include *k* disconnected subgraphs, each representing one species. In practice, the subgraphs could be linked by repeat sequences and some contigs are isolated from the main graph due to the sequencing errors, unbiased coverage, etc, called as dead ends. The performance of label propagation heavily relies on the number of edges and label density in the graph. The short contigs would be significantly affected by dead ends in two ways: (i) contigs would not be labeled if the dead end contains no label before propagation (Figure 1 dead end 1); (ii) labelling errors are induced if only a small number of contigs are labeled in the dead end (Figure 1 dead end 2).

**Figure 1.**
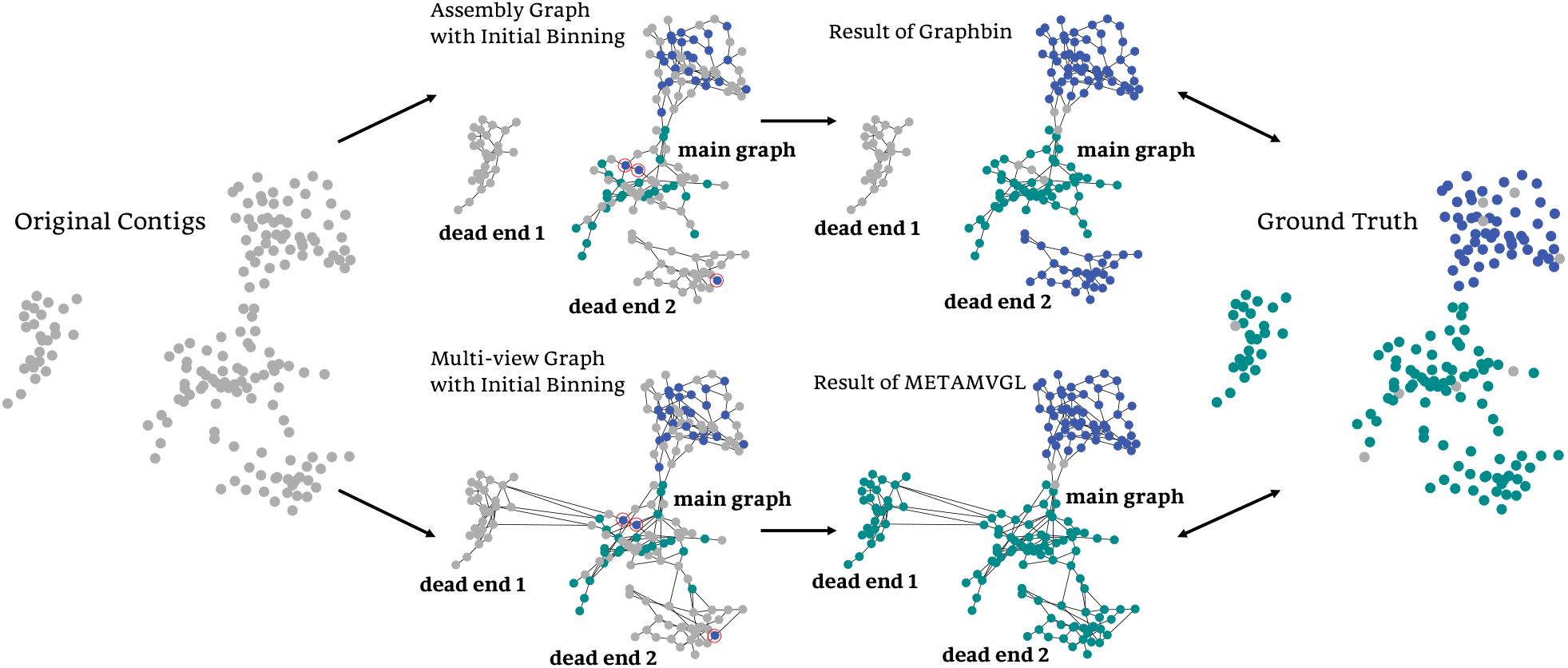
Visualization of the running process of METAMVGL compared with Graphbin in a simulated data. METAMVGL connected dead end 1 and 2 to the main graph by paired-end reads, also enhanced its connectivity. We observed (i) Graphbin failed to correct the two blue labels in the central of the graph, because it could not remove them before propagation due to lack of connectivity; (ii) Graphbin mislabeled all the vertices in dead end 2, caused by a small number of wrongly labeled vertices in the dead end; (iii) METAMVGL rescued all the labels in dead end 1 but Graphbin did not.

Here we present METAMVGL (Figure 2), a multiview graph-based metagenomic contig binning algorithm to address the above mentioned issues. METAMVGL not only considers the contig links from assembly graph but also involves the paired-end (PE) graph, representing the shared paired-end reads between two contigs. The two graphs are aggregated together by auto-weighting, where the weights together with the predicted labels are updated in a uniform framework (Nie et al. (2016), **Methods**). Figure 1 gives a proof-of-concept example on a simulated data, where the paired-end reads connect the two dead ends (dead end 1 and dead end 2) to the main graph. Our experiment results indicate METAMVGL substantially improves the binning performance of state-of-the-art binning algorithms, MaxBin2, MetaBAT2, MyCC, CONCOCT, SolidBin and Graphbin in all simulated, mock and *Sharon* datasets (Figure 3, Figure 4, Figure S1-S4). Comparing with assembly graph, PE graph can add up to 8942.37% vertices and 15114.06% edges to the main graph (Table S2 *Sharon* with MEGAHIT).

**Figure 2.**
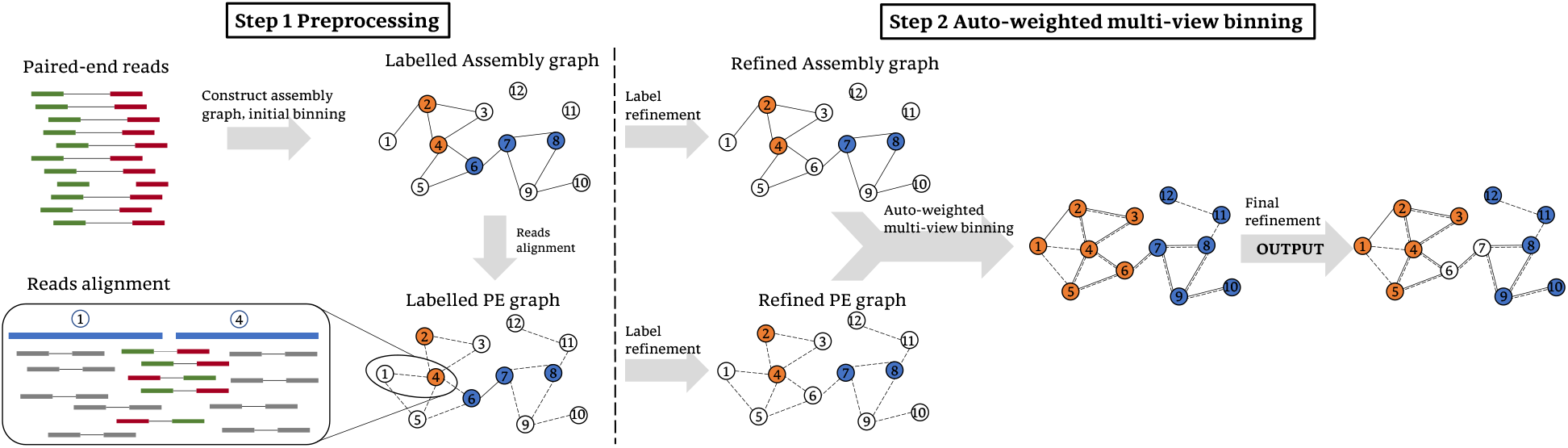
The workflow of METAMVGL. In step 1, we construct the assembly graph and PE graph by aligning paired-end reads to the contigs. The seed contigs are labeled by the available binning algorithms (vertices in orange and blue). In step 2, the ambiguous labels are removed if their neighbours are derived from different binning groups. METAMVGL applies auto-weighted multi-view graph-based algorithm to optimize the weight for each graph and predict binning groups of the unlabeled vertices. Finally, the second round of ambiguous labels removal is performed.

**Figure 3.**
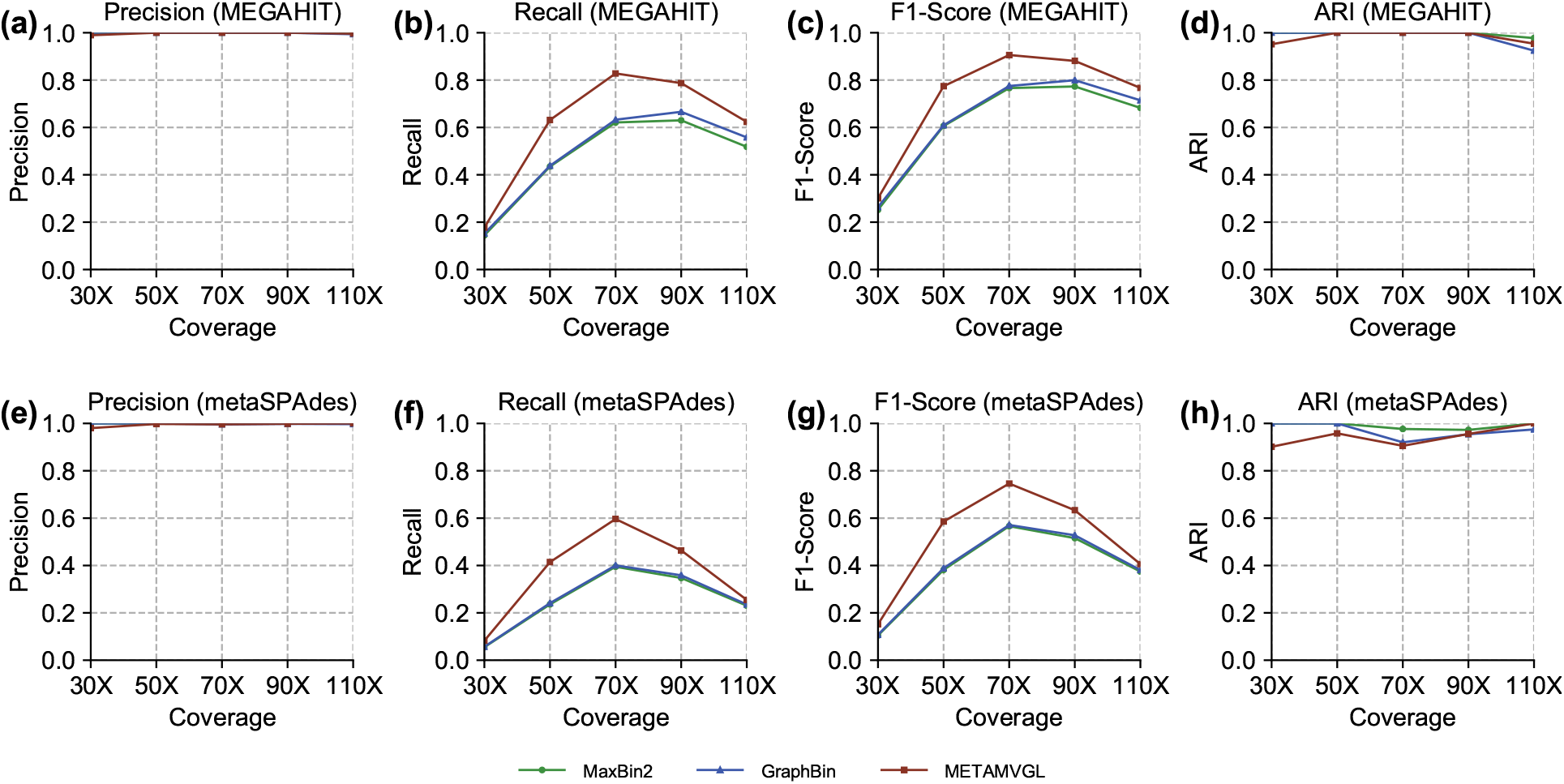
The evaluation results of METAMVGL, Graphbin with initial binning tool of MaxBin2 on simulated datasets. (a)-(g) are results based on MEGAHIT assembler, and (e)-(h) are results on metaSPAdes.

**Figure 4.**
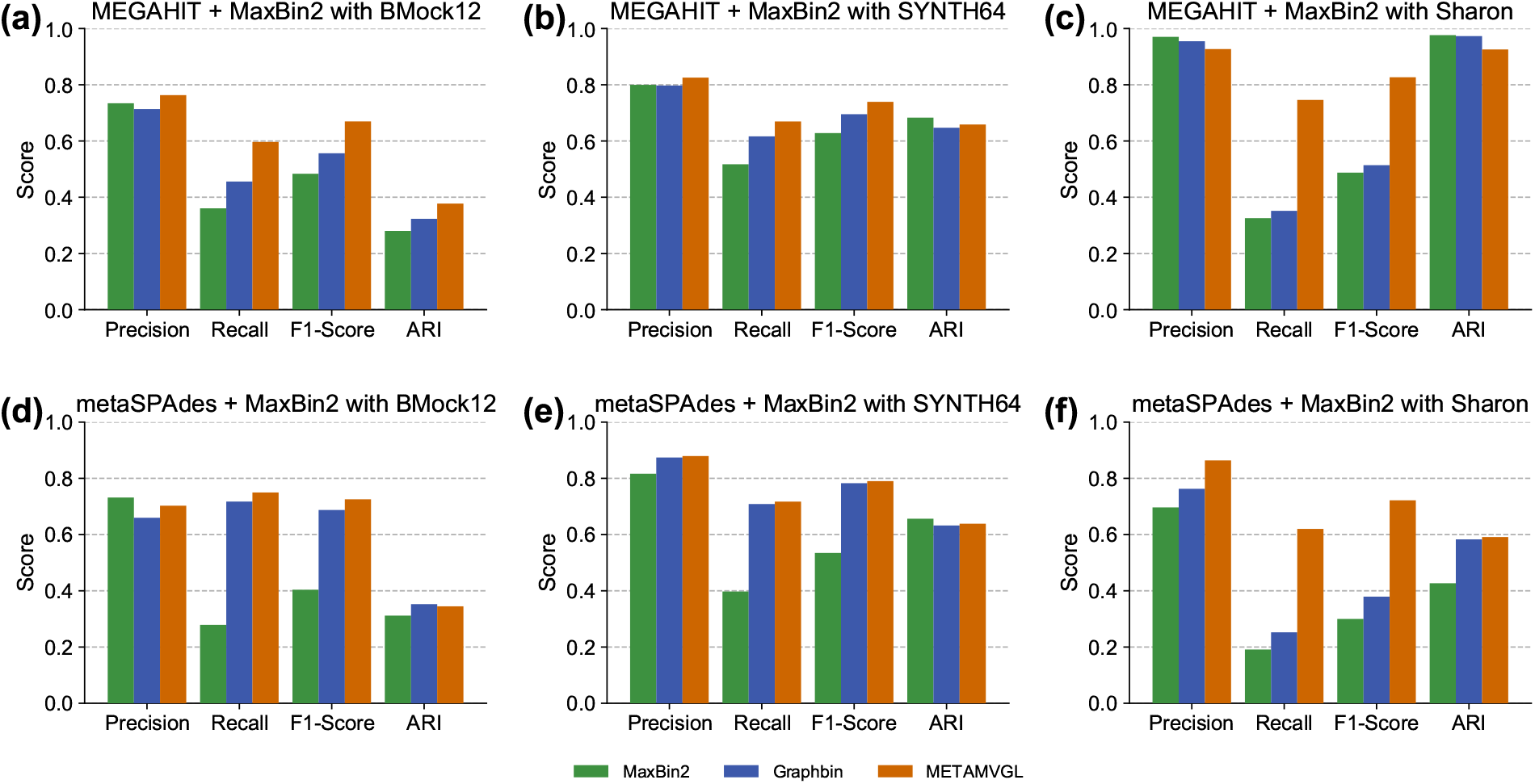
The results of METAMVGL, Graphbin with the initial binning tool of MaxBin2 in *BMock12, SYNTH64* and *Sharon*. MEGAHIT and metaSPAdes are used to generate assembly graphs. (a)-(b), (d)-(e) are results of mock datasets. (c) and (f) are *Sharon* results. Results for other initial binning tools (MetaBAT2, MyCC, CONCOCT and SolidBin) are in Figure S1-S4.

## 2 Materials and methods

Figure 2 illustrates the workflow of METAMVGL, which consists of two steps. In step 1, METAMVGL constructs the assembly graph and PE graph with contig labels computed by the available binning tools. In step 2, we remove the ambiguous labels of vertices if their neighbours are derived from different binning groups. The two graphs are merged by updating their weights iteratively. The unlabeled vertices are further predicted by label propagation. Finally, METAMVGL removes the ambiguous labels and generates the final binning results.

### 2.1 Step 1: Preprocessing

#### 2.1.1 Construct assembly graph

We define the assembly graph as *AG*(*V, E*), where the vertices *V* = {*v*_4_, *v*_2_, …,*v_n_*} represent contigs, and edge *e_i,j_* ∈ *E* exists if *v_i_* and *v_j_* are connected in the assembly graph and with *k* − 1 mer (continuous nucleotide of length *k* − 1) overlap. In principle, the assembly graph should include *k* unconnected subgraphs, each representing one species and we can easily recognize contig binning groups. In practice, the subgraphs could be linked due to the inter-species repeat sequences and complected by sequencing errors and unbalanced genomic coverage. Commonly the assembly graph includes one main graph and several dead ends. Figure 2 demonstrates an assembly graph with dead ends (vertices 11 and 12).

METAMVGL uses the assembly graph from metaSPAdes (Nurk et al. (2017)) and MEGAHIT (Li et al. (2015)). The original assembly graph of metaSPAdes is unitig-based graph, where the vertices are unitigs. The contigs are sets of unitigs after resolving short repeats. Hence we converted the unitig-based graph to contig-based graph by adding an edge *e_ij_*, if at least one of the unitigs belonging to *v_i_* and *v_j_* connected each other directly. MEGAHIT didn’t generate the assembly graph explicitly, so we used contig2fastg module of megahit_toolkit to generate the graph in fastg format.

#### 2.1.2 Construct PE graph

In order to deal with the dead ends in assembly graph, we constructed PE graph by aligning short-reads to the contigs. Paired-end reads were aligned to contigs using BWA-MEM(Li (2013)). For every two contigs, we maintained a paired-end reads set (*RS*), including the read names where the forward and reverse reads were aligned to the two contigs, respectively. We calculated the library insert size IS based on the uniquely aligned paired-end reads from the same contigs. To alleviate the influence of chimeric reads, we linked *v_i_* and *v_j_* if at least half of the reads in *RS_i,j_* came from the two stretches of length IS in both of the contigs (Bishara et al. (2018)). We denote the PE graph as *PE*(*V,E*), where *V* represents contigs, and E are edges supported by the PE links. Based on our observation, PE graph is complementary to the assembly graph to some degree, because edges in assembly graph only capture the overlaps between contigs, while PE graph can rescue the contig links due to the gaps. Figure 2 illustrates how dead ends of assembly graph can be linked to the main graph using PE graph.

#### 2.1.3 Initial binning

METAMVGL can accept the results from any contig binning algorithms to generate initial labels. In the experiments, we have tested MaxBin2, MetaBAT2, MyCC, CONCOCT and SolidBin in SolidBin-SFS mode. We used the default parameters for these algorithms except MetaBAT2, where its minimum contig length was set to 1.5kb to generate more labels.

### 2.2 Step 2: Auto-weighted multi-view binning

In auto-weighted multi-view binning, METAMVGL applies a label propagation based algorithm (Nie et al. (2016)) to learn the weights of assembly and PE graphs automatically and predict the unlabeled contigs in a uniformed framework. The binning is further purified by ambiguous labels removal.

#### 2.2.1 Remove ambiguous labels

The initial binning labels could be wrong especially for the contigs from repeat sequences and their influence would be amplified in label propagation. METAMVGL computes the distance between two vertices as the length of shortest path between them. Let *CLV*(*v*) be a set of labels from *v*’s closest labeled neighbours in graph *G* and contig *v*’s label is ambiguous if *CLV*(*v*) contains a label that is different from *v*’s label. *V_A_*(*G*) is a set including all the contigs with ambiguous labels in graph *G*. In Figure 2, the closest labeled vertices of *v*_6_ are {*v*_4_, *v*_7_}. Because *v*_4_ and *v*_6_ have different labels, the label of *v*_6_ is marked ambiguous. Algorithm 1 shows the process to remove ambiguous labels, where the function *CLV*(*G_i_*, *L*, *v_j_*) returns the indexes of closest labeled vertices of *v_j_* in graph *G_i_* (*i* ∈ 1, 2). *G*_1_ and *G*_2_ refer to assembly graph and PE graph, respectively. We applied Algorithm 1 to the graphs before and after label propagation.

**Algo 1:**
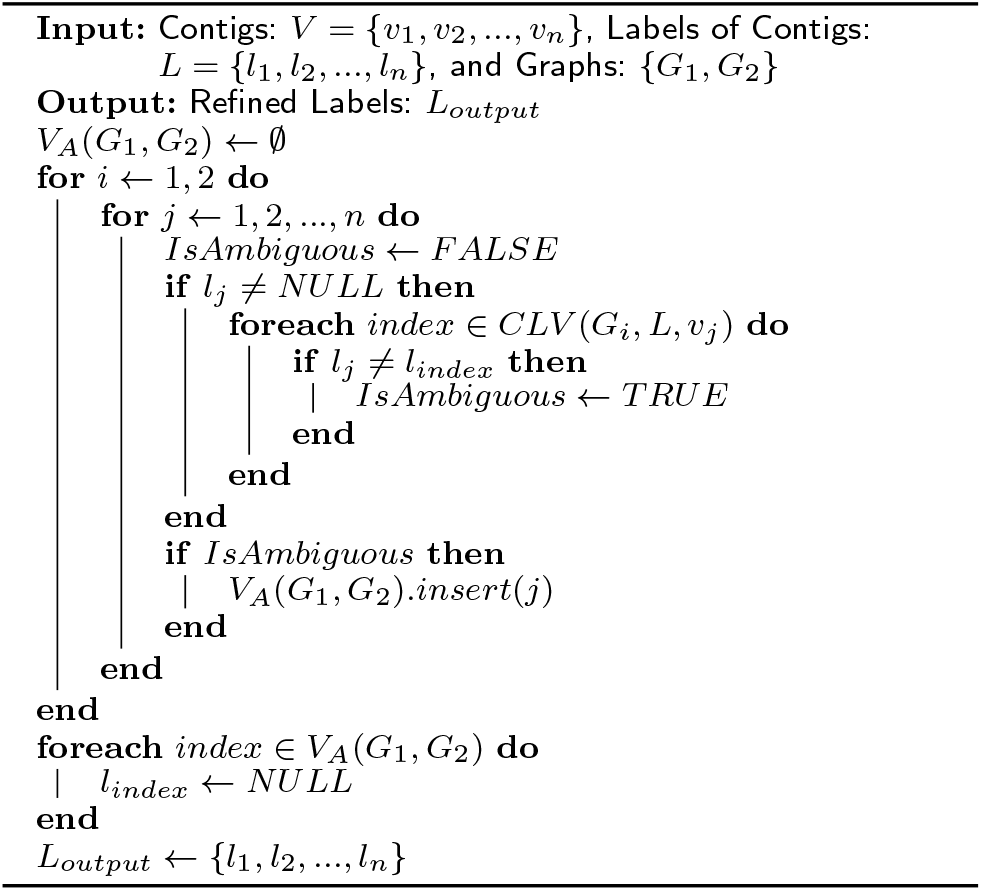
Remove Ambiguous Labels.

#### 2.2.2 Auto-weighted multi-view binning algorithm

Assume the initial binning algorithm annotates *l* contigs with c labels, denoted as 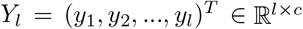, where *y_ij_* ∈ {0,1}, and *y_ij_* = 1 indicates the vertex *v_i_* is labeled species *j*. We define a matrix 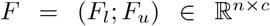, where *F_l_* = *Y_l_* and *F_u_* = (*f*_*l*+1_, *f*_*l*+2_,…,*f_n_*)^*T*^ are labels to be inferred. Let 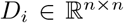 denote the degree matrix of *G_i_*, and 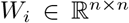 be the adjacent matrix of *G_i_*. The normalized Laplacian matrix of *G_i_* is defined as 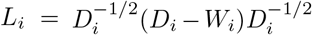. According to Nie et al. (2016), the above problem can be modeled as the following optimization problem:

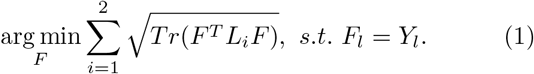

The optimization problem is converted to

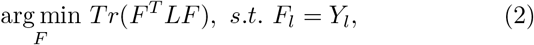

where 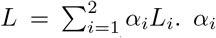 is the weight of *G_i_*, with initial values of 1/2. We partition L from (*l* + 1)^*th*^ row and column into four blocks as (*L_ll_, L_lu_; L_ul_, L_uu_*). *F* and *α_i_* can be updated alternatively until convergence by the following equations:

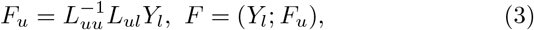

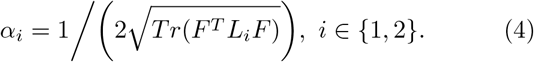

Equation 3 can be considered label propagation in the merged graph with automatically updated weight *α_i_*, hence *α_i_* implies the confidence of each graph. We infer the labels of all the vertices by *l_i_* = arg max_*j*_ *F_ij_, i* = 1,2,…,*n*. Algorithm 2 shows the process of auto-weighted multi-view binning, and Figure 2 is an example of this algorithm.

### 2.3 Datasets

#### 2.3.1 Simulated datasets

We simulated the metagenomic sequencing of three microbes with low, medium and high abundances from ATCC MSA-1003. The components are:

- ATCC_17978: 0.18%,
- ATCC_BAA-611: 1.8%,
- ATCC_700610: 18%.

**Algo 2:**
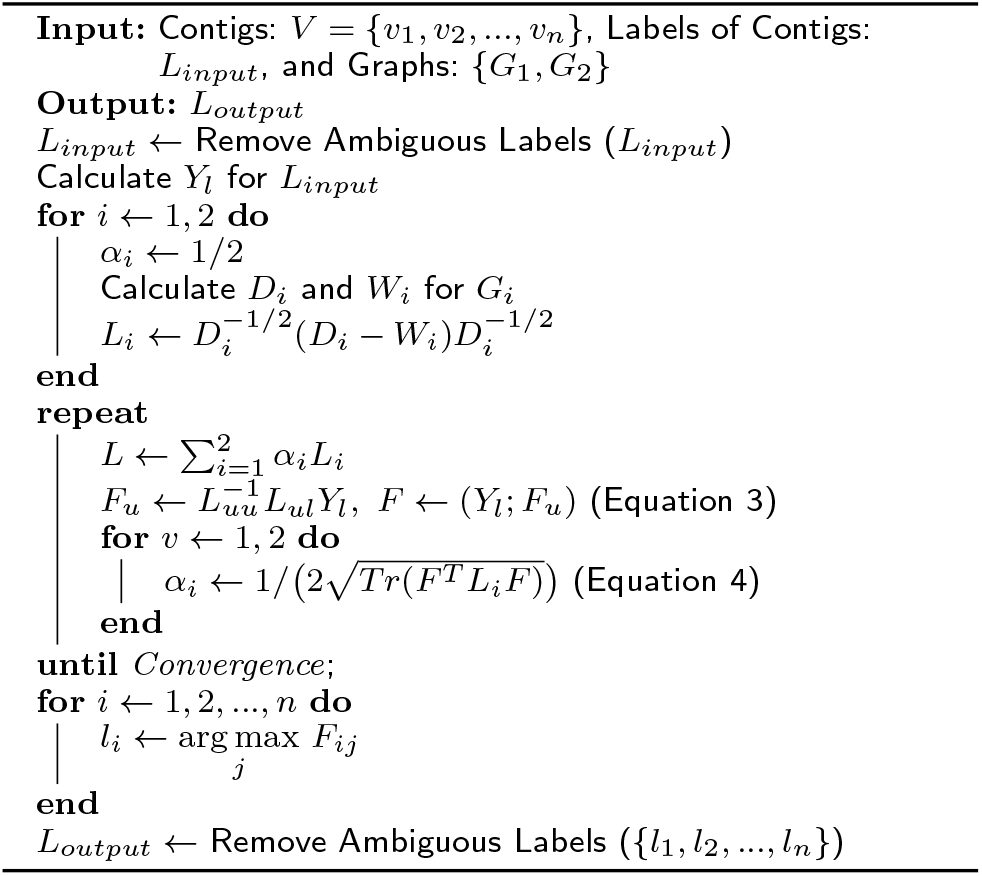
Auto-weighted multi-view binning.

We downloaded the complete reference genomes of the three species from the NCBI Nucleotide Database. CAMISIM (Fritz et al. (2019)) generated the shortreads for a mixture of the three genomes with corresponding abundance. Five simulated datasets were generated with read depths as 30x (ATCC_30x), 50x (ATCC_50x), 70x (ATCC_70x), 90x (ATCC_90x) and 110x (ATCC_110x).

#### 2.3.2 Mock datasets

We evaluated binning performance on the two mock community datasets following:

- *BMock12* refers to metagenomic sequencing for a mock community of 12 bacterial strains sequenced in Illumina HiSeq 2500 (Sevim et al. (2019); NCBI acc. no. SRX4901583). It contains 426.8 million 150bp reads with a total size of 64.4Gb.
- *SYNTH64* is metagenomic sequencing for a synthetic community with 64 diverse bacterial and archaea species (Shakya et al. (2013); NCBI acc. no. SRX200676), sequenced by Illumina HiSeq 2000 with read length 100bp and total size 5.8Gb.

#### 2.3.3 Real dataset

*Sharon* dataset (Sharon et al. (2013); NCBI acc. no. SRX144807) contains 18 metagenomic data from timeseries infant facial samples, sequenced by Illumina HiSeq 2000 with a total of 274.4 million 100bp reads. We combined all the 18 datasets for co-assembly and referred them as *Sharon*.

### 2.4 Evaluation criteria

To assess the binning result, we annotated the potential species the contigs come from. For simulated and mock datasets, we aligned the contigs to the available reference genomes and selected the ones with unique alignments. For *Sharon* dataset, we used kraken2 (Wood et al. (2019)) to annotate the species by k-mer similarities.

Assume there are *s* species, and the binning result have k groups. To evaluate the binning result, we define the assessment matrix 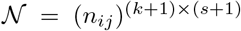, where *n_ij_* represents the number of contigs in *i^th^* bin that are annotated *j^th^* ground truth species. The (*k* + 1)^th^ row denotes unbinned contigs. The (*s* + 1)^th^ column indicates contigs without annotation. We applied (i) Precision, (ii) Recall, (iii) F1-Score and (iv) Adjusted Rand Index (ARI) to evaluate the binning performance. Let 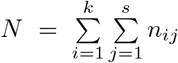, the four metrics were calculated as follows:

i. 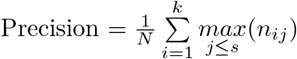,
ii. 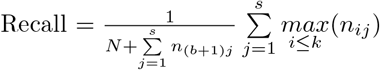,
iii. 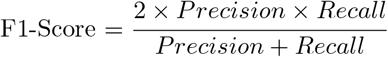,
iv. 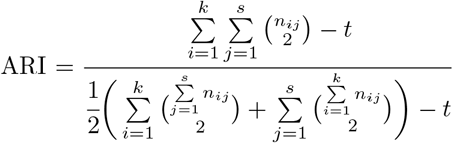,

where 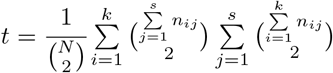.

## 3 Results

METAMVGL was compared with 6 binning tools, MaxBin2, MetaBAT2, MyCC, CONCOCT, SolidBin in SolidBin-SFS mode, and Graphbin based on the assembly graphs generated by MEGAHIT and metaSPAdes. We analyzed the binning performance on 5 simulated datasets with different sequence depths, 2 mock datasets of *BMock12* and *SYNTH64*, and a real community of *Sharon* dataset.

### 3.1 Evaluation on simulated datasets

Figure 3 shows the binning results of the simulated datasets. The contigs and assembly graph were generated by MEGAHIT (Figure 3 (a)-(d)) and metaSPAdes (Figure 3 (e)-(h)), MaxBin2 was applied as the initial binning tool for Graphbin and METAMVGL.

All the three binning tools (MaxBin2, Graphbin and METAMVGL) yielded extremely high precision and ARI (Figure 3 (a), (d), (e) and (h)), due to a low complexity of the simulated community. Because of considering assembly and PE graph jointly, METAMVGL labeled more contigs than Graphbin and MaxBin2 across different sequence depths, as shown in Figure 3 (b) and (d). We also found both Recall and F1-Score increased as sequencing depths became deeper until ATCC_70x (Figure 3 (b), (c), (f) and (g)). This observation was mirrored with the results in the paper of CAMISIM (Fritz et al. (2019)), suggesting a deep sequencing depth would introduce assembly noise even it might help in detecting the low-abundance microbes.

### 3.2 Evaluation on mock datasets

We illustrate the binning performance in mock datasets with initial binning tool of MaxBin2 (Figure 4 (a), (b), (d) and (e)). In general, the graph-based methods were always better than MaxBin2, but the results were different in choosing different assemblers. Graphbin and METAMVGL could significantly improve the recalls using metaSPAdes but the elevation became unobvious by MEGAHIT. This is because metaSPAdes generated more accurate and complete assembly graph than MEGAHIT. In the mock datasets, METAMVGL was just slightly better than Graphbin, suggesting the PE graph largely overlapped with the assembly graph (Table S2 *BMock12* and *SYNTH64*). This observation only occurred if perfect assembly graph was generated due to a low microbial complexity in the community.

Results of other initial binning tools can be found in Figure S1-S4, which are are akin to the observations from MaxBin2.

### 3.3 Evaluation on *Sharon* datasets

Figure 4 (c) and (f) describe the binning results in *Sharon* dataset. METAMVGL substantially increased the recall in comparison with both Graphbin (2.12 times for MEGAHIT; 2.46 times for metaSPAdes) and MaxBin2 (2.29 times for MEGAHIT; 3.24 times for metaSPAdes). METAMVGL also obtained the highest precision for metaSPAdes. All the three binning tools were comparable with ARI.

The outstanding recall of METAMVGL confirmed the capability of PE graph to connect dead ends to main graph when the assembly graph became incomplete in a real microbial community. We observed that MEGAHIT produced very fragmented assembly graph in *Sharon* dataset, in which the main graph only had 59 vertices with 64 edges, while the total number was 15,660 vertices (Table S2 *Sharon* with MEGAHIT). The fragmented assembly graph in real datasets was also mentioned as a limitation in Graphbin (Mallawaarachchi et al. (2020)). With PE graph, METAMVGL yielded 5,335 vertices and 9,737 edges in the main graph, rescuing a large number of unlabeled contigs from dead ends (Figure 4 (c)). Although the assembly graph was less fragmented (23.69% vertices in main graph) for metaSPAdes, PE graph still added 28.97% edges to the main graph (Table S2 *Sharon* with metaSPAdes), and improved the recall substantially.

## 4 Discussion

*De novo* assembly together with contig binning methods provided us a practical way to explore the novel microbes from metagenomic sequencing. But the current binning tools worked stably on only long contigs, the smaller ones were commonly neglected in the subsequent analysis. We observed a large proportion of contigs were shorter than 1kb, which introduced significant influence in bin completeness. Recent work (Mallawaarachchi et al. (2020)) proved short contigs could be rescued from assembly graph by considering their connections with the labeled ones. Assembly graph is accurate but relies heavily on the nature of microbial community. The high sequencing depth, sequencing errors and unbalanced coverage could generate considerable dead ends, which could introduce both missing labels and labelling errors (Figure 1).

In this paper, we developed METAMVGL, a multiview graph-based approach integrating both assembly and PE graphs. The model could automatically calculate the weights of the two graph and perform label propagation to predict the labels of short contigs. For the experiments, we observed METAMVGL could substantially increase the recalls with satisfying precision, especially for the sequencing data from the real complex microbial community. The integration of assembly and PE graph was still far from complete and there still require to consider the other information to reveal contig long range connectedness from long-read sequencing (PacBio and Oxford Nanopore) or linked-read sequencing (Tell-seq and stLFR).

## Additional files

**Additional file 1: Table S1.** Basic statics of contigs in real, mock, and simulated datasets, assembled by metaSPAdes and MEGAHIT.

**Additional file 2: Table S2.** Statistics of the biggest component of assembly graph, PE graph, and merged graph, in real, mock, and simulated datasets.

**Additional file 3: Figure S1.** The results of METAMVGL, Graphbin with initial binning tool of MetaBAT2 in *BMock12, SYNTH64* and *Sharon*.

**Additional file 4: Figure S2.** The results of METAMVGL, Graphbin with initial binning tool of MyCC in *BMock12, SYNTH64* and *Sharon*.

**Additional file 5: Figure S3.** The results of METAMVGL, Graphbin with initial binning tool of CONCOCT in *BMock12, SYNTH64* and *Sharon*.

**Additional file 6: Figure S4.** The results of METAMVGL, Graphbin with initial binning tool of SolidBin in *BMock12, SYNTH64* and *Sharon*.

## Acknowledgements

We thank Research Committee of Hong Kong Baptist University and Interdisciplinary Research Clusters Matching Scheme for their kindly support this project.

## Funding

LZ is supported by General Research Fund No. 22201419 HKSRA, IRCMS No. IRCMS/19-20/D02 HKBU, Guangdong Basic and Applied Basic Research Foundation, No. 2019A1515011046.

## Availability of data and materials

The source code of METAMVGL is publicly available at https://github.com/ZhangZhenmiao/METAMVGL. The Illumina short-reads of *BMock12, SYNTH64* and *Sharon* data are available in NCBI Sequence Read Archive (SRA), the accession numbers are SRX4901583, SRX200676 and SRX144807, respectively.

## Ethics approval and consent to participate

Not applicable.

## Competing interests

The authors have no conflicts of interest.

## Consent for publication

Not applicable.

## Authors’ contributions

LZ and ZMZ conceived the study. ZMZ implemented METAMVGL. ZMZ and LZ analyzed the results. ZMZ and LZ wrote the paper.

## Notes

### Competing Interest Statement

The authors have declared no competing interest.

