## Additional file 1 for "METAMVGL: a multi-view graph-based metagenomic contig binning algorithm by integrating assembly and paired-end graphs"

| **Dataset** | **Assembler** | | | **Contig Length** | | | | | | **N50** |
| --- | --- | --- | --- | --- | --- | --- | --- | --- | --- | --- |
|  |  |  |  | **>= 0 kb** | **>= 1kb** | **>= 5kb** | **>= 10kb** | **>= 25kb** | **>= 50kb** |  |
| **Sharon** | **MEGAHIT** | **#Contigs** | 15,660 | | 3,916 | 1,037 | 685 | 335 | 157 | 31,838 |
|  |  | **Total**  **Sequence (bp)** | 42,171,457 | | 36,660,321 | 31,010,065 | 28,493,777 | 22,864,853 | 16,533,264 |  |
|  | **metaSPAdes** | **#Contigs** | 42,249 | | 4,416 | 1,232 | 724 | 285 | 109 | 16,882 |
|  |  | **Total**  **Sequence (bp)** | 45,299,337 | | 36,421,145 | 29,953,620 | 26,367,231 | 19,526,067 | 13,439,978 |  |
| **SYNTH64** | **MEGAHIT** | **#Contigs** | 20,645 | | 13,322 | 5,891 | 3,585 | 1,745 | 857 | 53,957 |
|  |  | **Total**  **Sequence (bp)** | 202,393,964 | | 198,278,962 | 180,676,140 | 164,407,087 | 135,828,139 | 104,750,298 |  |
|  | **metaSPAdes** | **#Contigs** | 28,245 | | 12,979 | 5,398 | 3,202 | 1,591 | 869 |  |
|  |  | **Total**  **Sequence (bp)** | 202,047,505 | | 196,651,034 | 178,405,210 | 162,978,840 | 137,854,369 | 112,375,186 | 63,744 |
| **BMock12** | **MEGAHIT** | **#Contigs** | 3,820 | | 1,583 | 815 | 626 | 423 | 264 | 115,661 |
|  |  | **Total**  **Sequence (bp)** | 49,966,643 | | 48,937,816 | 47,320,490 | 45,974,215 | 42,728,338 | 37,050,583 |  |
|  | **metaSPAdes** | **#Contigs** | 5,805 | | 1,244 | 723 | 544 | 363 | 226 | 156,751 |
|  |  | **Total**  **Sequence (bp)** | 49,587,091 | | 48,622,707 | 47,464,913 | 46,172,793 | 43,152,535 | 38,286,155 |  |
| **ATCC**  **_110x** | **MEGAHIT** | **#Contigs** | 1,354 | | 455 | 184 | 67 | 16 | 13 | 13,302 |
|  |  | **Total**  **Sequence (bp)** | 4,524,180 | | 4,122,710 | 3,414,050 | 2,603,810 | 1,886,016 | 1,766,038 |  |
|  | **metaSPAdes** | **#Contigs** | 4,069 | | 341 | 173 | 77 | 20 | 13 | 13,800 |
|  |  | **Total**  **Sequence (bp)** | 5,112,009 | | 4,077,400 | 3,626,933 | 2,942,216 | 2,118,241 | 1,891,318 |  |
| **ATCC**  **_90x** | **MEGAHIT** | **#Contigs** | 1,462 | | 628 | 132 | 34 | 15 | 13 | 9,235 |
|  |  | **Total**  **Sequence (bp)** | 4,497,985 | | 4,098,612 | 2,851,373 | 2,201,991 | 1,934,911 | 1,842,766 |  |
|  | **metaSPAdes** | **#Contigs** | 3,747 | | 545 | 140 | 33 | 14 | 13 | 7,536 |
|  |  | **Total**  **Sequence (bp)** | 4,891,842 | | 3,981,713 | 2,892,030 | 2,168,292 | 1,904,152 | 1,876,457 |  |
| **ATCC**  **_70x** | **MEGAHIT** | **#Contigs** | 1,867 | | 832 | 56 | 17 | 14 | 11 | 5,205 |
|  |  | **Total**  **Sequence (bp)** | 4,344,725 | | 3,795,379 | 2,200,625 | 1,967,420 | 1,903,433 | 1,782,161 |  |
|  | **metaSPAdes** | **#Contigs** | 4,016 | | 768 | 46 | 19 | 14 | 13 | 3,991 |
|  |  | **Total**  **Sequence (bp)** | 4,704,608 | | 3,673,586 | 2,161,151 | 1,992,797 | 1,904,153 | 1,876,458 |  |
| **ATCC**  **_50x** | **MEGAHIT** | **#Contigs** | 2,231 | | 701 | 22 | 19 | 16 | 12 | 5,249 |
|  |  | **Total**  **Sequence (bp)** | 3,981,388 | | 3,114,900 | 1,991,621 | 1,974,704 | 1,910,007 | 1,749,509 |  |
|  | **metaSPAdes** | **#Contigs** | 4,905 | | 582 | 19 | 19 | 14 | 13 | 2,089 |
|  |  | **Total**  **Sequence (bp)** | 4,442,198 | | 2,884,357 | 1,983,220 | 1,983,220 | 1,891,349 | 1,863,654 |  |
| **ATCC**  **_30x** | **MEGAHIT** | **#Contigs** | 1,856 | | 183 | 20 | 18 | 15 | 12 | 115,029 |
|  |  | **Total**  **Sequence (bp)** | 3,102,130 | | 2,207,747 | 1,988,327 | 1,976,456 | 1,912,318 | 1,801,699 |  |
|  | **metaSPAdes** | **#Contigs** | 5,019 | | 134 | 18 | 18 | 14 | 13 | 27,695 |
|  |  | **Total**  **Sequence (bp)** | 3,758,397 | | 2,132,037 | 1,982,280 | 1,982,280 | 1,904,162 | 1,876,467 |  |

Table S1. Basic statics of contigs in real, mock, and simulated datasets, assembled by metaSPAdes and MEGAHIT.
