## Additional file 2 for "METAMVGL: a multi-view graph-based metagenomic contig binning algorithm by integrating assembly and paired-end graphs"

| **Dataset** | **Assembler** | **All Vertices** | **Assembly Graph** | | | **PE Graph** | | | | **Merged Graph** | | |
| --- | --- | --- | --- | --- | --- | --- | --- | --- | --- | --- | --- | --- |
|  |  |  | **#Vertices** | **#Edges** | | **#Vertices** | | **#Edges** | | **#Vertices** | **#Edges** | |
| **Sharon** | **MEGAHIT** | 15660 | 59 | | 64 | | 4967 | | 8947 | 5335 | | 9737 |
|  | **metaSPAdes** | 42249 | 10009 | | 14668 | | 5780 | | 9738 | 10273 | | 18918 |
| **SYNTH64** | **MEGAHIT** | 20645 | 577 | | 686 | | 7641 | | 10186 | 8221 | | 11343 |
|  | **metaSPAdes** | 28245 | 15065 | | 25423 | | 8475 | | 12227 | 16736 | | 32780 |
| **BMock12** | **MEGAHIT** | 3820 | 310 | | 331 | | 1068 | | 1650 | 1118 | | 1725 |
|  | **metaSPAdes** | 5805 | 4049 | | 6910 | | 1712 | | 3054 | 4093 | | 8177 |
| **ATCC_110x** | **MEGAHIT** | 1354 | 30 | | 30 | | 145 | | 177 | 196 | | 233 |
|  | **metaSPAdes** | 4069 | 53 | | 90 | | 241 | | 282 | 271 | | 381 |
| **ATCC_90x** | **MEGAHIT** | 1462 | 24 | | 26 | | 109 | | 132 | 127 | | 157 |
|  | **metaSPAdes** | 3747 | 51 | | 87 | | 74 | | 104 | 89 | | 101 |
| **ATCC_70x** | **MEGAHIT** | 1867 | 17 | | 19 | | 76 | | 90 | 110 | | 131 |
|  | **metaSPAdes** | 4016 | 49 | | 86 | | 83 | | 106 | 88 | | 156 |
| **ATCC_50x** | **MEGAHIT** | 2231 | 15 | | 16 | | 54 | | 66 | 56 | | 72 |
|  | **metaSPAdes** | 4905 | 42 | | 77 | | 38 | | 53 | 44 | | 95 |
| **ATCC_30x** | **MEGAHIT** | 1856 | 14 | | 14 | | 46 | | 54 | 62 | | 78 |
|  | **metaSPAdes** | 5019 | 44 | | 78 | | 45 | | 60 | 52 | | 102 |

Table S2. Statistics of the biggest component of assembly graph, PE graph, and merged graph, in real, mock, and simulated datasets.
