## Supplementary figures and images for "METAMVGL: a multi-view graph-based metagenomic contig binning algorithm by integrating assembly and paired-end graphs"

### Additional file 3

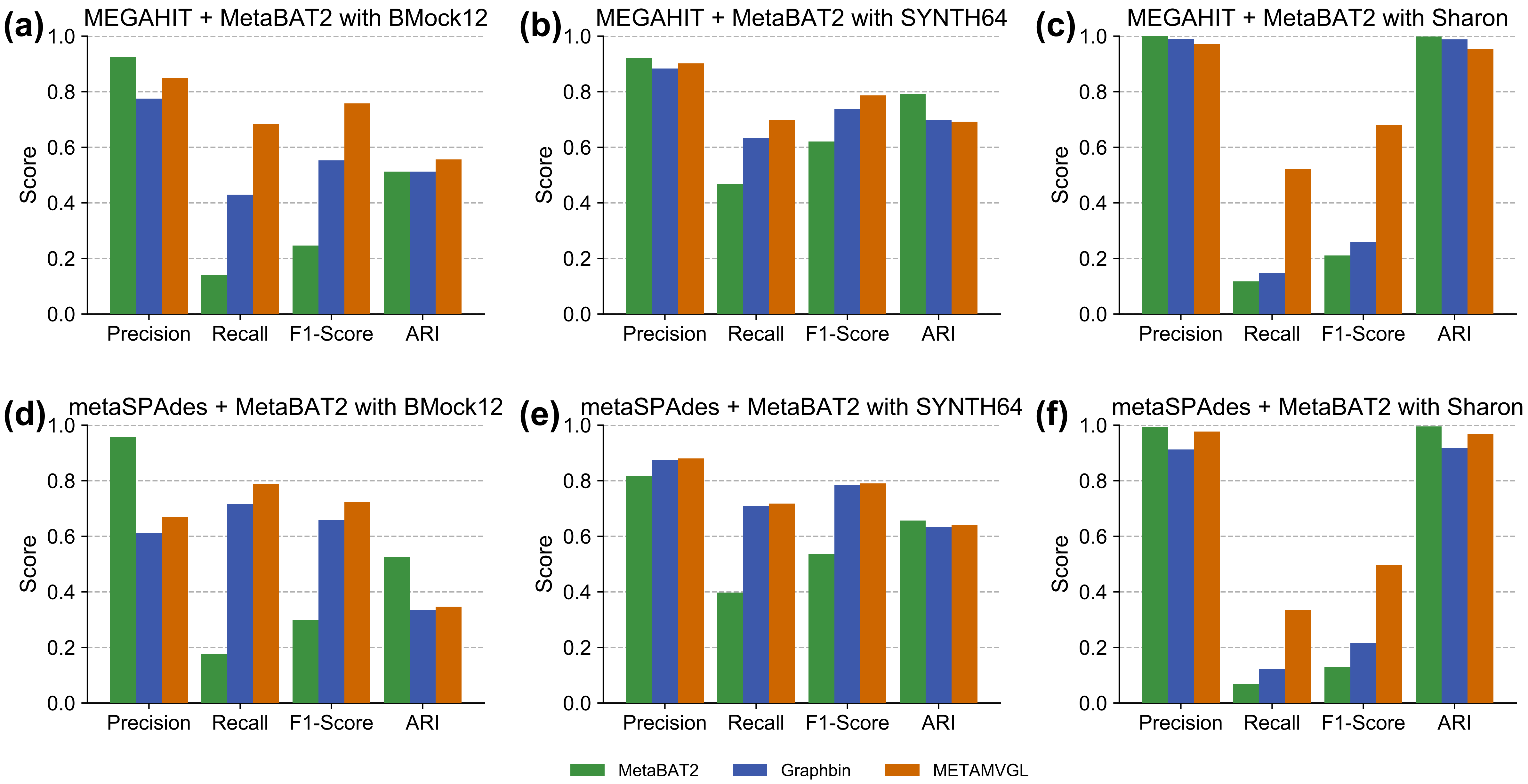

### Additional file 4

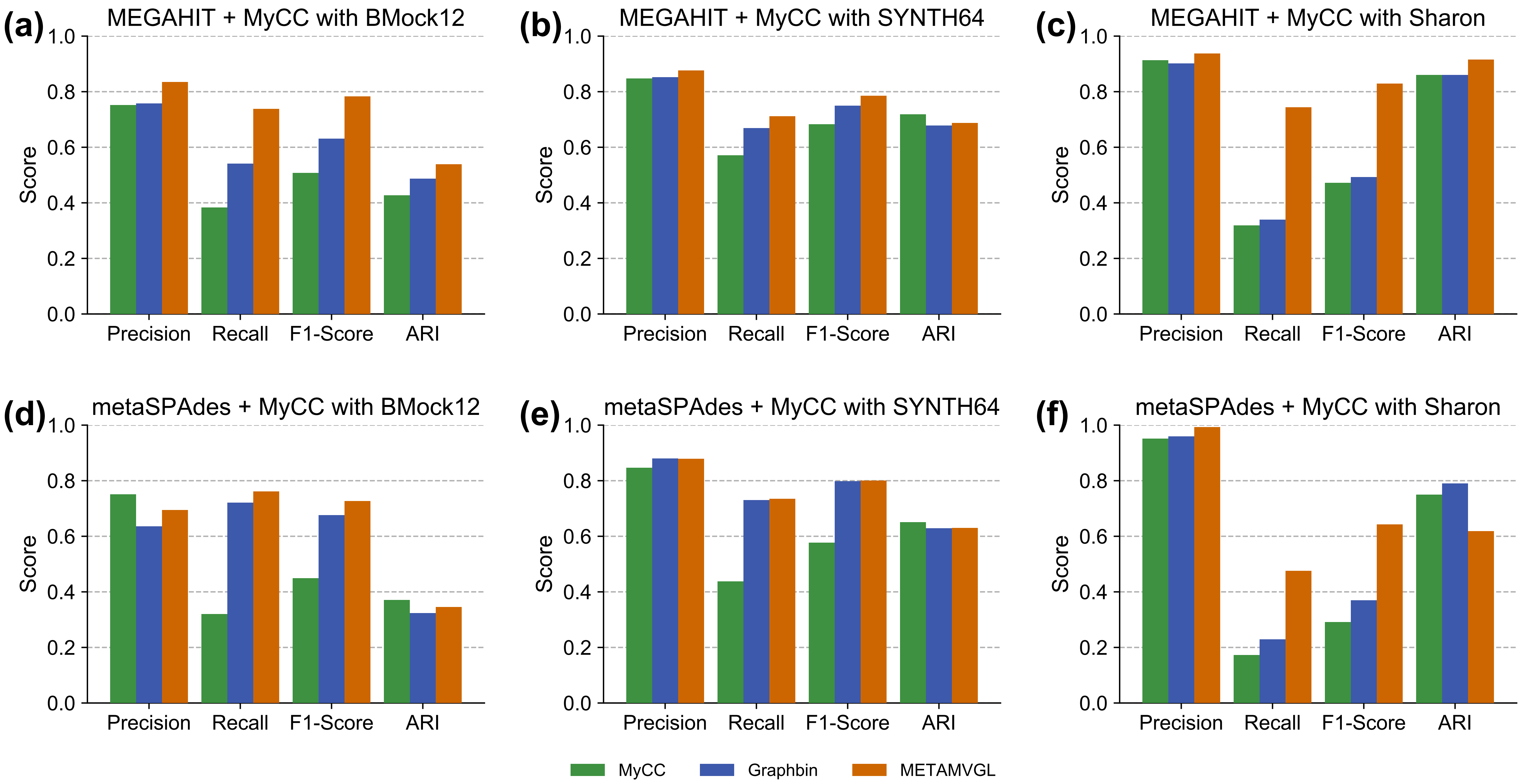

### Additional file 5

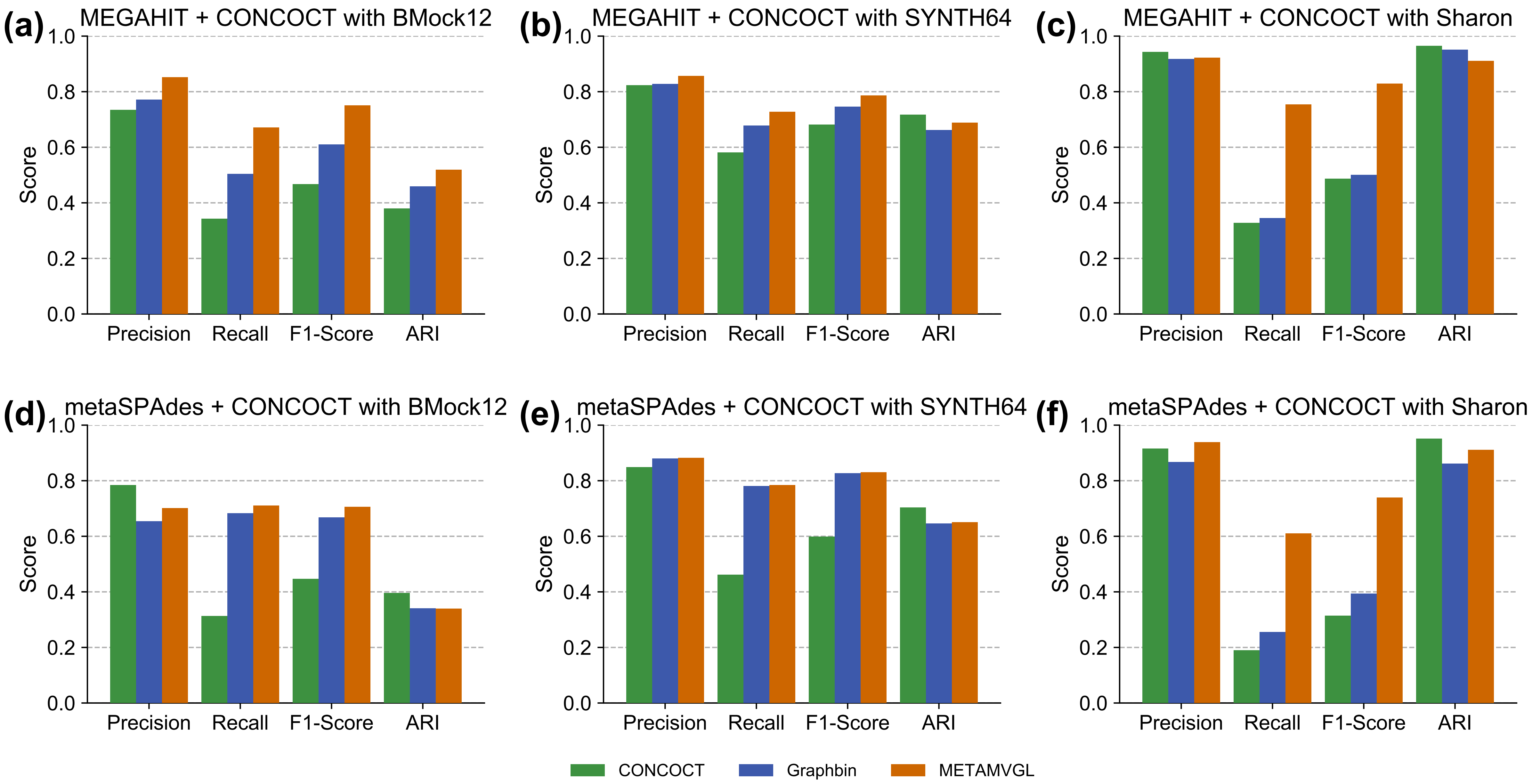

### Additional file 6

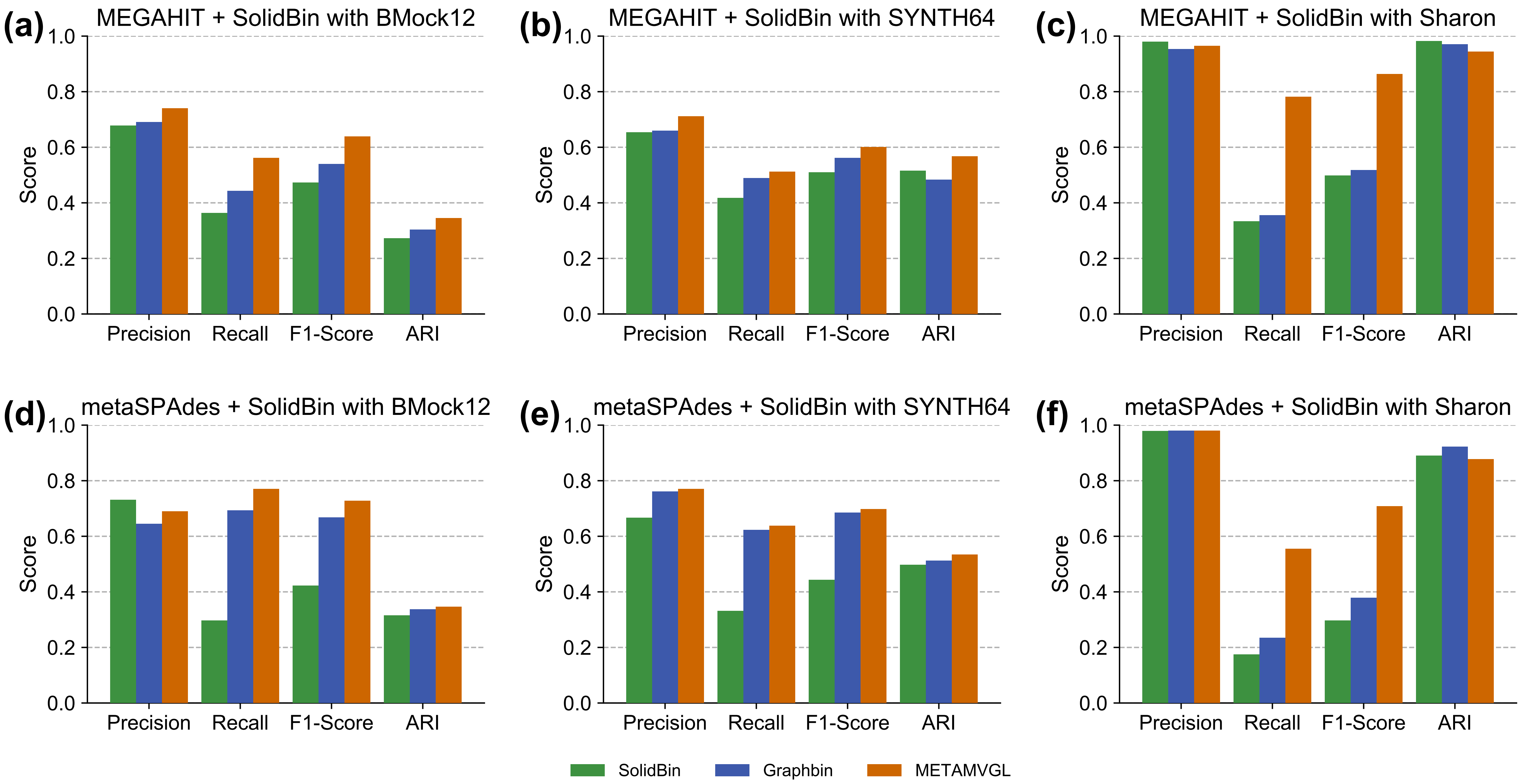
